# How prior and p-value heuristics are used when interpreting data

**DOI:** 10.1101/2023.09.03.556128

**Authors:** Ethan Hermer, Ashley A. Irwin, Dominique G. Roche, Roslyn Dakin

## Abstract

Scientific conclusions are based on the ways that researchers interpret data, a process that is shaped by psychological and cultural factors. When researchers use shortcuts known as heuristics to interpret data, it can sometimes lead to errors. To test the use of heuristics, we surveyed 623 researchers in biology and asked them to interpret scatterplots that showed ambiguous relationships, altering only the labels on the graphs. Our manipulations tested the use of two heuristics based on major statistical frameworks: (1) the strong prior heuristic, where a relationship is viewed as stronger if it is expected *a priori*, following Bayesian statistics, and (2) the p-value heuristic, where a relationship is viewed as stronger if it is associated with a small p-value, following null hypothesis statistical testing. Our results show that both the strong prior and p-value heuristics are common. Surprisingly, the strong prior heuristic was more prevalent among inexperienced researchers, whereas its effect was diminished among the most experienced biologists in our survey. By contrast, we find that p-values cause researchers at all levels to report that an ambiguous graph shows a strong result. Together, these results suggest that experience in the sciences may diminish a researcher’s Bayesian intuitions, while reinforcing the use of p-values as a shortcut for effect size. Reform to data science training in STEM could help reduce researchers’ reliance on error-prone heuristics.

**Significance Statement:** Scientific researchers must interpret data and statistical tests to draw conclusions. When researchers use shortcuts known as heuristics, it can sometimes lead to errors. To test how this occurs, we asked biologists to interpret graphs that showed an ambiguous relationship between two variables, and report whether the relationship was strong, weak, or absent. We altered features of the graph to test whether prior expectations or a statistic called the p-value could influence their interpretations. Our results indicate that both prior expectations and p-values can increase the probability that researchers will report that ambiguous data shows a strong result. These findings suggest that current training and research practices promote the use of error-prone shortcuts in decision-making.

## INTRODUCTION

In science, researchers frequently interpret data visualizations to draw inferences. These interpretations are made when researchers act as producers of new knowledge, and when they act as consumers and reviewers of results presented by other researchers. When data interpretation decisions and their underlying criteria are clear, they can aid scientific reproducibility and consensus (Parker et al. 2016). On the other hand, conclusions based on small datasets and ambiguous relationships are frequently unreliable and can differ among researchers (Ioannidis 2019) (Bakker et al. 2019). When faced with this type of ambiguous data, what drives researchers’ interpretations?

One strategy that researchers may use are heuristics, or simple rules that speed up the interpretation process (Kahneman and Tversky 1972, Gigerenzer 2004, Gigerenzer and Gaissmaier 2011). These shortcuts are efficient, but they can also be error prone (Gigerenzer and Gaissmaier 2011). Here, we test whether researchers use two simple heuristics when interpreting ambiguous data. As the subjects for this study, we focused on biologists, because statistical inference is so important in the life sciences and can be a source of error (Weissgerber et al. 2016). Our goal was to test whether heuristics increase the probability that researchers report that ambiguous data shows a strong result.

We tested two heuristics derived from two major statistical frameworks: the strong prior heuristic, and the p-value heuristic. First, we define the strong prior heuristic as interpreting a relationship as stronger if that relationship is expected to occur *a priori*. This is related to a Bayesian statistical framework. In Bayesian statistics, evidence is formalized by updating a prior expectation with information from observed data. All else being equal, a prior belief that a particular relationship is strong will increase the posterior certainty that it is strong, as computed via Bayes’ theorem. Some aspects of the human decision-making process have been hypothesized to follow this process (Ma et al. 2006, Bornstein et al. 2017, de Lange et al. 2018). For example, when interpreting the strength of a relationship presented in a scientific paper, researchers will interpret the results of the paper in the context of their knowledge of other previous findings, which may reflect an intuitive application of a Bayesian framework. However, strong priors can also be a potential source of bias and error. For example, when a researcher has a favoured outcome or pet hypothesis, they might develop corresponding “wishful” priors that shape their interpretation of data.

We define the p-value heuristic as interpreting a simple relationship as stronger if the relationship is associated with a very small p-value. This heuristic is based on null-hypothesis statistical testing (NHST), a statistical framework that has been widely taught in statistics courses since the 1950s (e.g., Chavalarias et al. 2016). The p-value is intended to measure the probability of obtaining a given result, or range of results, under the assumption that a specific null hypothesis is true (Wasserstein and Lazar 2016). NHST introduces a threshold method, whereby a p-value below a given threshold (typically 0.05) leads to rejection of the null hypothesis, and the interpretation that there is evidence for the current hypothesis (Amrhein et al. 2017, Benjamin et al. 2017). There are many ways that NHST and p-values are misused and misinterpreted that been reviewed widely in recent decades (e.g., Gelman 2016, Greenland et al. 2016, Wasserstein and Lazar 2016, Lakens 2021). Yet this recognition has not dampened the widespread use of p-values, both in statistical education and research practices (e.g., Chavalarias et al., 2016). Although p-values are not measures of effect size, they are often misinterpreted as such (e.g., Amrhein et al., 2017; Goodman, 2008; Wasserstein and Lazar, 2016). Thus, when faced with an ambiguous dataset but a small p-value, the prominence of this heuristic may cause researchers to conclude that the effect size is inappropriately large or strong.

To test the use of these two heuristics, we surveyed researchers in biology and asked them to interpret the strength of relationships shown in scatterplots (Figure 1). The scatterplots showed bivariate relationships that were weakly positive, but ambiguous. Each participant was asked to interpret two scatterplots in series. The first interpretation task was designed to test the use of the strong prior heuristic, with two treatments that altered the scatterplot axis labels, as shown in Figure 1B. Participants assigned the “prior” treatment saw labels representing two variables, heart disease and smoking, that would be commonly expected to be positively correlated *a priori* (e.g., McCoy et al. 1992). Participants assigned the “no prior” treatment saw labels representing internet browsing and height, two variables that are not expected to be correlated.

**Figure 1.**
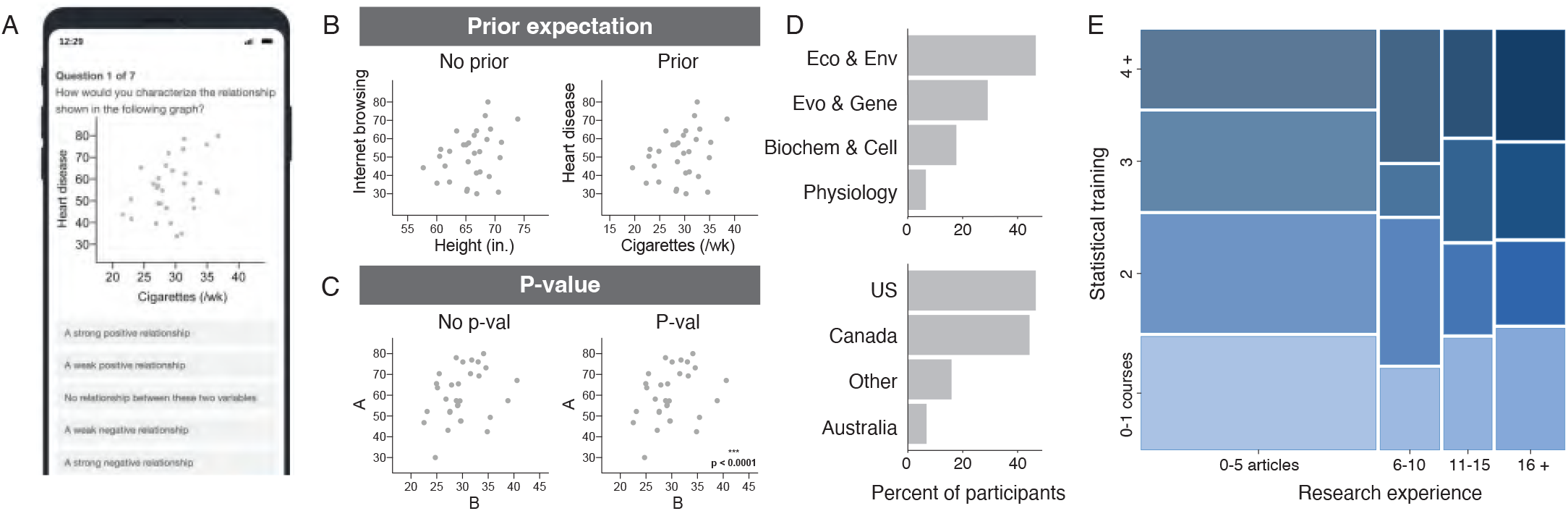
Testing the use of data interpretation heuristics by biologists. We surveyed researchers in biology to determine their use of two data interpretation heuristics: strong priors, and p-values. (A) Participants were asked to interpret scatterplots that showed a positive, but ambiguous (uncertain) relationships. (B-C) Treatments used in the heuristic tests. (B) The first question tested the influence of priors by manipulating axis labels. Note that the treatment on the left right (internet use and height) represents two variables that are not expected to be related, whereas the treatment on the right (heart disease and smoking) represents a relationship with a strong positive prior. (C) The second question tested the influence of a very small p-value, “p < 0.0001”. Note that the true p-values for the scatterplots ranged from 0.07 – 0.11, and “p < 0.0001” was a mismatch to the data shown. For each test within (B) and (C), participants were randomly assigned to one treatment or the other using a between-subjects design. Scatterplots were also randomly assigned from a set of options shown in Figure S1. (D) Bar graphs showing the geographic and disciplinary distribution of biologists who participated in the study (n = 623). (E) We also assessed participants’ research experience and statistical training. We defined experience as the number of peer reviewed papers that participant had authored in the last five years, and training as the number of statistical courses they had taken or taught. The mosaic plot in (E) illustrates the makeup of these characteristics in the participant sample.

The second interpretation task was designed to test the use of the p-value heuristic, with two treatments that altered the presence of a very small p-value in the lower right corner of the scatterplot, as shown in Figure 1C. Importantly, the “p-value” treatment showed a p-value that was unrelated to the data in the scatterplot. This recreates a scenario where a statistic is incorrectly presented alongside data, which can be a source of many real research errors. The participants were researchers in biology that we recruited from professional societies and university departments (Figure 1D). Participants were randomly assigned to one treatment group for each task in a between-subjects design, with the assignment performed independently for each of the two interpretation tasks.

We hypothesized that both the strong prior heuristic and the p-value heuristic would be used to interpret ambiguous data. A key question is how data interpretation biases are also shaped by STEM education and training (Weissgerber et al. 2016). We thus determined participants’ level of training by asking them how many statistics courses they had taken or taught. We also assessed participants’ level of research experience based on how many peer-reviewed research articles they had published within the last five years (Figure 1E). If training or experience reinforces Bayesian intuitions, then one might expect the strong prior heuristic to be more prevalent among researchers who have taken or taught more statistics courses, or published more papers. An alternative possibility is that training and/or experience may cause researchers to disregard their prior expectations (e.g., to maintain objectivity), and reduce the prevalence of the strong prior heuristic.

When considering the p-value heuristic, we expected that higher levels of training or experience would reduce the use of the p-value heuristic, either because researchers learn through personal experience (trial-and-error) that p-values are not reliable indicators of an effect size, or because they have had more exposure to academic and pedagogical discussion on the misuse of p-values in biology (Greenland et al. 2016). Alternatively, the p-value heuristic may be entrenched among biological researchers, regardless of their level of experience or training.

## RESULTS AND DISCUSSION

Our experiment shows that both types of heuristics, the strong prior and the p-value heuristic, are commonly used by biologists (Figure 2). In the absence of priors, the ambiguous data were reported as “strong” about 10% of the time (Figure 2A). But when the labels on the graphs corresponded to prior expectations, researchers were about twice as likely to report a strong relationship (Figure S2A, right). This prior-induced shift in interpretations was almost entirely driven by the least experienced participants in our sample. Examining the p-value, this second heuristic had a similar effect: the presence of a small p-value nearly doubled the probability that a researcher would report a strong relationship (Figure S2B, right). Because we used a between-subjects design, we cannot tell whether any one researcher was using one (or both) heuristics; nevertheless, our results show that expert interpretations can easily be nudged by error-prone shortcuts.

**Figure 2.**
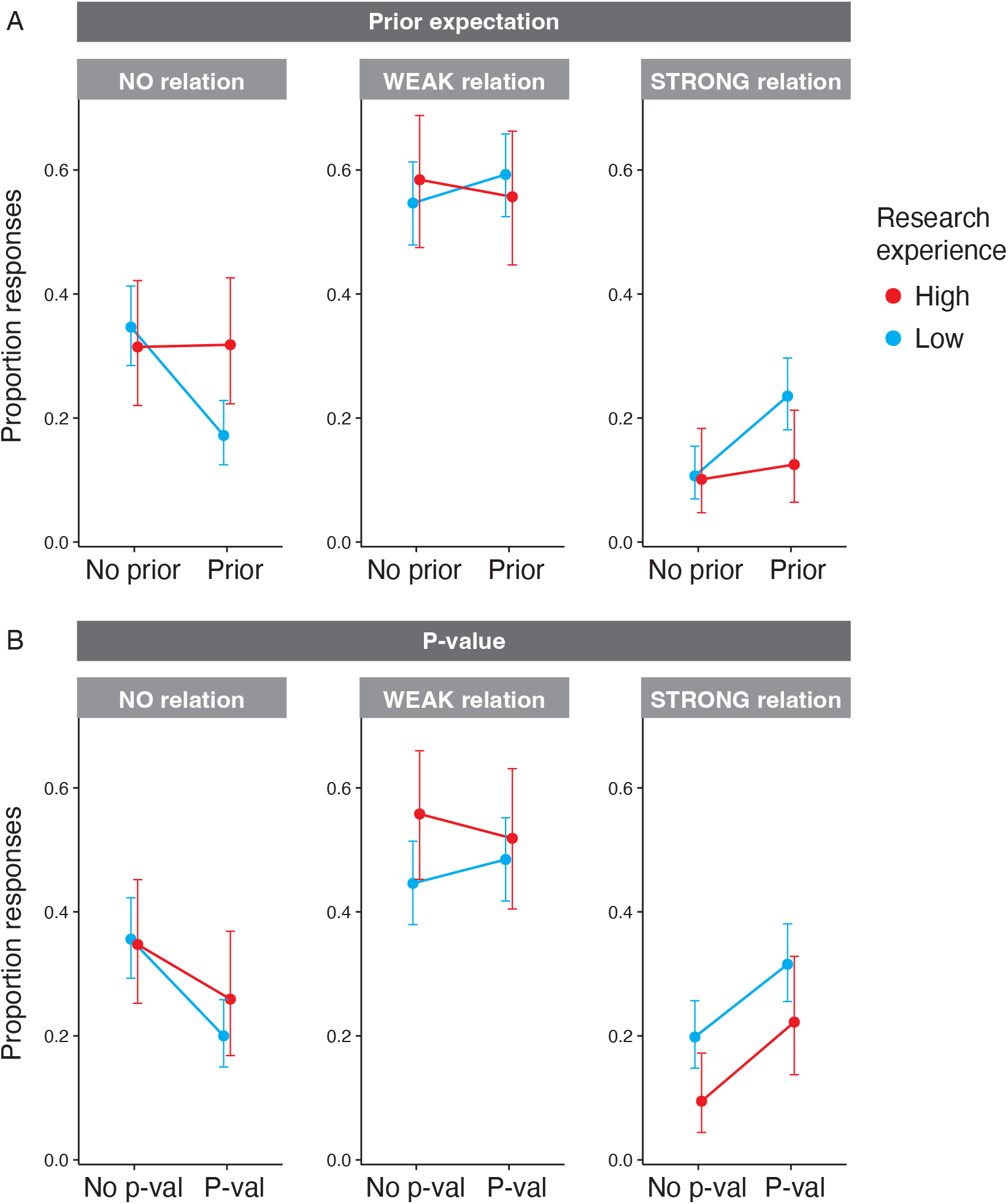
The use of prior and p-value heuristics in biology. Effect of (A) prior expectations and (B) small p-values on how researchers interpreted ambiguous data. Points show the proportion of participants who reported “No relationship” (left column), “A weak positive relationship” (middle), or “A strong positive relationship” (right). Error bars show 95% confidence intervals, and the connecting lines within each panel are used to highlight the treatment effects. (A) In the first test, we find early-stage researchers with relatively low experience (blue) are influenced by strong priors, because the prior treatment increased the likelihood that they would report a strong result. Yet prior expectations had little to no effect on the most experienced researchers in our survey (red). (B) In the second test, we find that a label with a small p-value increased the likelihood that researchers would report a strong result, regardless of their level of experience or training. Note that experience was scored using four levels in the analysis; it was collapsed into two categories here for visualization purposes only. See Figures S2 and S3 for additional details.

The influence of priors depended on a participant’s level of research experience: the prior manipulation had the strongest effect on researchers in our survey with the fewest recent papers (Figure 2A, Tables S2 and S4). By contrast, the seasoned researchers who had the greatest number of recent papers were not detectably influenced by the prior manipulation (Figure 2A). Interestingly, we did not detect any modulating effect of experience or training on the use of the p-value heuristic (Figure 2B, Tables S3 and S5), suggesting that researchers at all levels use p-values as a shortcut to interpret relationship strength.

The fact that experienced researchers rely less on priors raised the further possibility that skilled researchers may be more sensitive to the data at hand. If this hypothesis is correct, then we would expect the interpretations of more experienced researchers to depend more strongly on the exact scatterplot viewed, whereas the interpretations of inexperienced researchers should be less sensitive to scatterplot variations. Given that the participants in our survey viewed six different scatterplots (Figure S1), we chose to explore this idea with two additional *post-hoc* analyses. The first additional step we took was to check whether response variation, across the two interpretation tasks pooled, could be explained by an interaction between the scatterplot viewed and the participants’ experience level. We found some evidence for this dependency (Figure S4A; delta AICc = 2.25 in favour of a model with the experience x scatterplot interaction; LRT p = 0.03). Second, we estimated the proportion of response variation explained by scatterplot variants, for experienced and inexperienced participants, respectively (Figure S4B). Among experienced researchers, scatterplot differences explained roughly 12% of the response variation (R^2^glmm 95% CI = 0–24%), whereas scatterplot differences explained only 4% of the response variation for researchers with the least experience (R^2^glmm 95% CI = 0–9%).

These results suggest that researchers with greater experience of the research process may learn to pay more heed to the data at hand, or that they may learn to become more skeptical of ambiguous results. Another (non-exclusive) possibility is that more experienced researchers may become less influenced by extraneous features on graphs. It is important to treat these additional analyses with caution because our study was not designed to test scatterplot differences *per se*. Indeed, when designing our experiment, we chose a set of scatterplots that were as similarly ambiguous as possible as shown in Figure S1. As such, additional experiments are needed test what mechanism may cause changes in the way researchers interpret data as they gain experience. Based on our results, we hypothesize that direct experience of replicating samples (and critically, failing to replicate your own ambiguous findings) may be a key stimulus that drives data interpretation behaviour.

Our results show that the p-value heuristic is widespread. Furthermore, our findings also suggest that p-value heuristics are insensitive to high levels of training and experience. This is striking, because problems associated with common use of p-values are not new (Wasserstein and Lazar 2016, Muff et al. 2022), and recently, critiques of p-values have been widely shared in broad-impact journals (Amrhein et al. 2019). In the statement by the American Statistical Association on the problems with p-values, a quote by George Cobb nicely summarizes the issue: “We teach it because it’s what we do; we do it because it’s what we teach” (ASA). Our results in this study confirm this view: the (mis)use of p-values is entrenched among biologists at all levels, even if most researchers are aware of these problems (Lakens 2021). We suggest, like Cobb, that this is a cultural problem, where an error-prone shortcut is maintained because of the convenience of heuristic thinking, combined with social transmission. We further suggest that heuristics may be a causal factor underlying many errors in research. While we only tested each heuristic in isolation here, it is easy to imagine how the combination of wishful priors *and* overinterpreted p-values could lead researchers to fool themselves and others.

Given the risk that heuristics pose for research reproducibility, an important question is how can STEM training and culture be changed to reduce heuristic thinking? Some authors have called for an elimination of p-values altogether, recommending that they be replaced by other frameworks, such as confidence intervals, Information Theoretic approaches, or Bayesian approaches, as examples (Amrhein et al. 2019, Halsey 2019, Muff et al. 2022). Yet, even in these other frameworks, users can still easily slip into heuristic or dichotomous thinking, for example, by applying NHST-based thresholds to Bayesian posterior distributions (Gelman 2016, Amrhein et al. 2019, Lakens 2021, Muff et al. 2022). Therefore, the solution may not be to just switch statistical frameworks, but shift towards teaching users how to think more slowly and critically about evidence and uncertainty (Berger and Berry 1988, Gelman 2016, Muff et al. 2022). A key contribution of our experiment here is that it provides a way for future studies to test how different reporting standards influence how data are viewed. It can also be used to trial the downstream effects of different pedagogical approaches to quantitative training in data science. Such research would be particularly valuable if paired with scholarship on teaching and learning, especially among students who will form the future research community.

## MATERIALS AND METHODS

### Survey

To test the use of heuristics, we developed a short online survey using the Qualtrics XM platform and distributed it to academic researchers in the biological sciences. The first page of the survey informed participants that the aim of the survey was to investigate how scientists interpret data visualizations. Screenshots of the survey are included in the supplement. Upon completion of the survey, participants were offered the option to enter a draw for a $40 gift card. All methods were approved by the Carleton University Research Ethics Board (CUREB-B #114980).

The first two survey questions tested the influence of the prior and p-value heuristics, respectively, by asking participants interpret a scatterplot showing 30 datapoints with an ambiguous, weak positive relationship (Figure 1A-C). Participants could choose from five options: “A strong positive relationship”, “A weak positive relationship”, “No relationship between these two variables”, “A weak negative relationship”, or “A strong negative relationship”. Although none of the scatterplots showed a negative relationship, these five options were included to represent all possibilities for such a relationship.

The first survey question tested the influence of the prior heuristic by varying the axis labels on the scatterplot. For the “prior” condition (right of Figure 1B), the axis labels indicated a measure of heart disease on the y-axis in relation to cigarette use on the x-axis, because the relationship between smoking and heart disease is widely known (McCoy et al. 1992) and expected from extensive previous research conducted over the past 50 years (Jacobs et al. 1999). For the “no prior” condition (left of Figure 1B), the axis labels indicated a measure of internet browsing on the y-axis in relation to height on the x-axis, because there is no general expectation that these two measures would be related.

The second survey question tested the influence of the p-value heuristic by varying the presence of a statistically significant p-value as shown in Figure 1C. For the “p-val” condition, a small p-value “p < 0.0001” was printed in the lower right corner of the scatterplot, paired with three asterisks (right of Figure 1C). In the “no p-val” condition, these features were absent (left of Figure 1C). In this p-value test, the axis labels were always uninformative (labels “A” and “B”, as shown in Figure 1C). Note that the true p-values for the data shown in the scatterplots ranged from 0.07 – 0.11, with Pearson’s correlation coefficients ranging from 0.29 – 0.24 (i.e., the data shown had a weak positive, but uncertain, relationship). As such, the “p-val” condition provided information that was incorrect for the data shown.

Treatments were randomly assigned in a between-subjects design, with the random assignment performed independently for each of the two test questions via the Qualtrics random presentation function. The scatterplots were also randomly assigned to participants from a set of options using the same method (see supplement for details).

Following the two data interpretation questions, we surveyed participants on their level of statistical training and research experience, their field(s) of study, their highest degree obtained, and the country of their primary university (see supplement for details). Most participants selected 1-2 biological fields of study from a list of 15 options (see supplement for details). Additional details on the biological research disciplines of participants are provided in the supplement.

We distributed the survey in two rounds. In January 2021, we sent an initial invitation to the membership of the Canadian Society of Zoologists, a professional organization of biologists (n = 139 participants completed the survey in this first round). In March 2021, we distributed the invitation to members of another professional organization, the Ecological Society of America, and to biology departments at several universities in the United States, Canada, Australia, and the United Kingdom (n = 526 participants completed the survey in this second round). Survey questions on the highest degree obtained and the country of primary university were only added in the second distribution. Based on those responses, approximately 70% of participants had completed a postgraduate degree, 17% were currently working on a postgraduate degree, and 7% had completed a bachelor’s or undergraduate as their highest degree (a further 6% did not answer this question).

## Data analysis

All data processing and analyses were performed in R 4.3.1 (R Core Team 2023). The median time to complete the survey was 2.1 minutes. We excluded 39 participants from further analysis because they did not answer one of the training and experience questions. For each heuristic tests, we excluded a further 3 responses that reported a negative relationship in the scatterplot, which we assume to be errors. After these exclusions, the sample size for each heuristic test was 623 survey participants.

We performed a separate statistical analysis for each heuristic test: first, the use of prior expectations, and second, the use of the p-value heuristic. We modelled each response as an ordinal categorical variable (no relationship, weak positive relationship, or strong positive relationship). We used the clm function in the package ordinal 2022.11-16 (Christensen 2022) to fit ordinal regression models examining the effect of the heuristic treatment. All models included predictors for treatment as well as the scatterplot dataset shown (see Figure S1). For each heuristic test, we fit five candidate models to examine the influence of researcher training or experience:

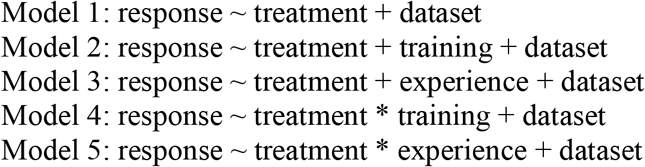

Both the training and the experience variables were scored as numerical variables ranging from 1 to 4. For training, a score of 1 was defined as the participant reporting having taken part in 0 or 1 statistics course(s), a score of 2 as 2 statistics courses, a score of 3 as 3 courses, and a score of 4 as 4 or more courses. We based our measure of experience on the extent of the participants’ reported recent experience publishing articles in the peer-reviewed literature over the past 5 years. A score of 1 was defined as the participant having published 0-5 articles, a score of 2 as 6-10 articles, a score of 3 as 11-15 articles, and a score of 4 as 16 or more articles over the past 5 years. The distribution of these variables is shown in Figure 1E. Models 4 and 5 included interactive effects of the treatment and the participants’ level of training or experience, to test whether the use of a particular heuristic depends on these characteristics.

We used the model.sel function in the package MuMIn 1.47.5 (Barton 2023) to rank model support based on the Akaike information criterion, AICc. In the first test of the prior heuristic, model 5 was best supported by a wide margin (delta AICc > 3.3; Akaike weight 0.7). In the second test of the p-value heuristic, support for the candidate models was not as well resolved and models 3, 1, and 5 were all within 1.1 AICc units of each other. We thus considered the simplest model of these three, model 1, as best-supported in that set; importantly, we also verified that all interpretations of the p-value heuristic test are unchanged when examining or averaging the other closely competing models.

We checked the proportional odds assumption for all ordinal regression model fits using the nominal_test function in the ordinal package (Christensen 2022). The proportional odds assumption was met in all models of the prior heuristic test, but not in all models of the p-value heuristic test. Therefore, as an additional step, we also examined the p-value heuristic test using a multinomial regression modelling framework. The multinomial regression model treats the responses (none, weak, strong) as three unordered categories. To do this, we used the multinom function in the package nnet 7.3-19 (Venables and Ripley 2002). We verified that the conclusions from the p-value heuristic test were unchanged when using either the ordinal or multinomial modelling frameworks, and present the ordinal model results in Figure 2.

After interpreting our main analyses in Figure 2, our results raised the possibility that more experienced researchers may be more sensitive to the data at hand when interpreting the strength of relationships depicted in scatterplots. If this hypothesis is correct, we would expect that differences among scatterplot datasets should explain more of the response variation among highly experienced researchers, as compared to less experienced researchers. To explore this possibility, we conducted two further *post-hoc* analyses shown in Figure S4. First, we pooled the participants’ responses across the two experiments, such that there were two responses per participant (i.e., repeated measures). We classified their responses as reporting that there was a positive relationship (either strong or weak) or no relationship at all. We also stratified the participants as either low research experience (0-5 recent articles, the most common category in our sample), or high research experience (6 or more recent articles). We then fit a generalized linear mixed-effects model (GLMM) with a binomial error distribution using the package lme4 1.1-34 (Bates et al. 2015) to explore the effect of research experience on sensitivity to scatterplot variants. The predictors included in the model were experience, scatterplot dataset, and the experience * dataset interaction. Participant identity was included in the model to account for repeated measures. See Figure S4A for details. This first *post-hoc* analysis indicated some evidence for an experience * dataset interaction (delta AICc = 2.25 over a model without the interaction), consistent with the hypothesis that highly experienced researchers are more sensitive to the data at hand when interpreting the strength of a scatterplot.

As a second *post-hoc* analysis, we also estimated the proportion of response variation that can be attributed to differences between scatterplots. To do this, we fit two further GLMMs for each researcher experience class (low and high experience), specifying scatterplot dataset as a random effect, and estimated a R^2^glmm using the MuMIn package. We obtained confidence intervals for the R^2^glmm estimates via parametric bootstrapping. These results are presented in Figure S4B. Consistent with the result described above, the R^2^glmm estimates suggest that dataset variants can explain slightly more of the response variation among highly experienced researchers (12%), as compared with less experienced researchers (4%). Although the effect of dataset variants was fairly subtle, we note that the six datasets in our study were chosen to have similarly high levels of ambiguity (as shown in Figure S1). Thus, further study examining a greater range of dataset variants would be useful to fully test how experience shapes a researchers’ sensitivity to the data at hand when interpreting graphs.

## References

Amrhein, V., S. Greenland, and B. McShane. 2019. Retire statistical significance. Nature 567:306–307.

Amrhein, V., F. Korner-Nievergelt, and T. Roth. 2017. The earth is flat (p > 0:05): Significance thresholds and the crisis of unreplicable research. PeerJ 2017.

Bakker, A., J. Cai, L. English, G. Kaiser, V. Mesa, and W. Van Dooren. 2019. Beyond small, medium, or large: points of consideration when interpreting effect sizes. Educational Studies in Mathematics 102:1–8.

Barton, K. 2023. MuMIn: Multi-Model Inference.

Bates, D., M. Mächler, B. Bolker, and S. Walker. 2015. Fitting linear mixed-effects models using lme4. Journal of Statistical Software 67.

Benjamin, D. J., J. O. Berger, M. Johannesson, B. A. Nosek, E.-J. Wagenmakers, R. Berk, K. A. Bollen, B. Brembs, L. Brown, C. Camerer, D. Cesarini, C. D. Chambers, M. Clyde, T. D. Cook, P. De Boeck, Z. Dienes, A. Dreber, K. Easwaran, C. Efferson, E. Fehr, F. Fidler, A. P. Field, M. Forster, E. I. George, R. Gonzalez, S. Goodman, E. Green, D. P. Green, A. G. Greenwald, J. D. Hadfield, L. V. Hedges, L. Held, T. Hua Ho, H. Hoijtink, D. J. Hruschka, K. Imai, G. Imbens, J. P. A. Ioannidis, M. Jeon, J. H. Jones, M. Kirchler, D. Laibson, J. List, R. Little, A. Lupia, E. Machery, S. E. Maxwell, M. McCarthy, D. A. Moore, S. L. Morgan, M. Munafó, S. Nakagawa, B. Nyhan, T. H. Parker, L. Pericchi, M. Perugini, J. Rouder, J. Rousseau, V. Savalei, F. D. Schönbrodt, T. Sellke, B. Sinclair, D. Tingley, T. Van Zandt, S. Vazire, D. J. Watts, C. Winship, R. L. Wolpert, Y. Xie, C. Young, J. Zinman, and V. E. Johnson. 2017. Redefine statistical significance. Nature Human Behaviour 2:6–10.

Berger, J. O., and D. A. Berry. 1988. Statistical analysis and the illusion of objectivity. American Scientist 76:159–165.

Bornstein, A. M., M. W. Khaw, D. Shohamy, and N. D. Daw. 2017. Reminders of past choices bias decisions for reward in humans. Nature Communications 8:15958.

Chavalarias, D., J. D. Wallach, A. H. T. Li, and J. P. A. Ioannidis. 2016. Evolution of reporting p values in the biomedical literature, 1990-2015. JAMA 315:1141.

Christensen, R. 2022. ordinal - Regression Models for Ordinal Data.

Gelman, A. 2016. The problems with p-values are not just with p-values. The American Statistician 70.

Gigerenzer, G. 2004. Mindless statistics. The Journal of Socio-Economics 33:587–606.

Gigerenzer, G., and W. Gaissmaier. 2011. Heuristic decision making. Annual Review of Psychology 62:451–482.

Goodman, S. 2008. A dirty dozen: twelve p-value misconceptions. Seminars in Hematology 45:135–140.

Greenland, S., S. J. Senn, K. J. Rothman, J. B. Carlin, C. Poole, S. N. Goodman, and D. G. Altman. 2016. Statistical tests, P values, confidence intervals, and power: a guide to misinterpretations. European Journal of Epidemiology 31:337–350.

Halsey, L. G. 2019. The reign of the p -value is over: what alternative analyses could we employ to fill the power vacuum? Biology Letters 15:20190174.

Ioannidis, J. P. A. 2019. What have we (not) learnt from millions of scientific papers with p values? The American Statistician 73:20–25.

Jacobs, D. R., H. Adachi, I. Mulder, D. Kromhout, A. Menotti, A. Nissinen, and H. Blackburn. 1999. Cigarette smoking and mortality risk: twenty-five–year follow-up of the seven countries study. Archives of Internal Medicine 159:733–740.

Kahneman, D., and A. Tversky. 1972. Subjective probability: A judgment of representativeness. Cognitive psychology 3:430–454.

Lakens, D. 2021. The practical alternative to the p value is the correctly used p value. Perspectives on Psychological Science. 16: 639–648.

de Lange, F. P., M. Heilbron, and P. Kok. 2018. How do expectations shape perception? Trends in Cognitive Sciences 22:764–779.

Ma, W. J., J. M. Beck, P. E. Latham, and A. Pouget. 2006. Bayesian inference with probabilistic population codes. Nature Neuroscience 9:1432–1438.

McCoy, S. B., F. X. Gibbons, T. J. Reis, M. Gerrard, C. A. E. Luus, and A. Von Wald Sufka.1992. Perceptions of smoking risk as a function of smoking status. Journal of Behavioral Medicine 15:469–488.

Muff, S., E. B. Nilsen, R. B. O’Hara, and C. R. Nater. 2022. Rewriting results sections in the language of evidence. Trends in Ecology and Evolution 37:203–210.

Parker, T. H., W. Forstmeier, J. Koricheva, F. Fidler, J. D. Hadfield, Y. E. Chee, C. D. Kelly, J. Gurevitch, and S. Nakagawa. 2016. Transparency in ecology and evolution: real problems, real solutions. Trends in Ecology & Evolution 31:711–719.

R Core Team. 2023. R: A language and environment for statistical computing. R Foundation for Statistical Computing, Vienna, Austria.

Venables, W., and B. Ripley. 2002. Modern Applied Statistics with S. Fourth. Springer, New York.

Wasserstein, R. L., and N. A. Lazar. 2016. The ASA’s statement on p-values: context, process, and purpose. American Statistician 70:129–133.

Weissgerber, T. L., V. D. Garovic, J. S. Milin-Lazovic, S. J. Winham, Z. Obradovic, J. P. Trzeciakowski, and N. M. Milic. 2016. Reinventing biostatistics education for basic scientists. PLOS Biology 14:e1002430.

