## supplement for "How prior and p-value heuristics are used when interpreting data"

To assess participants' level of statistical training, the survey asked: "Approximately how many courses in statistics have you completed (as a student) or taught (as an instructor)? (Include undergraduate and graduate level)". Responses were reported as an integer value from "0" up to "9 or more". Because most values fell in the low range, we categorized responses into four levels: a score of 1 was defined as the participant reporting 0 or 1 statistics course(s), a score of 2 as 2 courses, a score of 3 as 3 courses, and a score of 4 as 4 or more courses.

To assess participants' level of research experience, the survey asked: "How many peer-reviewed research articles have you published in the last 5 years?". Participants selected a category from the following options: "0-5", "6-10", "11-15", or "16 or more".

To summarize the main disciplinary affiliations of the participants, the survey asked participants to: "Please select one or more field(s) that you identify most closely with:" and they could select one or more of the major fields of biology listed in Table S1 below. The labels were derived from Web of Science research categories. Most participants selected 1-2 labels, but some selected 3 or more. We performed a network analysis of the chosen subfields using the package igraph 1.3.4 to link participants (as nodes) by the number of shared subfields (as weighted edges). We then used the walktrap community finding algorithm to identify the major disciplinary cliques or affiliation categories within the participant network. Table S1 below summarizes the results of this analysis, to help characterize the makeup of our participant sample.

**Table S1. Major affiliations of survey participants.** Participants could select one or more biological field(s) that they most closely identified with. These labels were taken from research categories used by Web of Science (n = 612 participants completed this question). The second column in the table shows the percent of participants who selected a given subfield on the survey question (note that the sum is greater than 100 because participants could choose as many as subfields as they wished). We used a network analysis of these data to group participants into four major affiliation communities: (i) ecology and environment, (ii) evolution and genetics, (iii) biochemistry and cell biology, and (iv) physiology. The table presents the percentage of participants in each affiliation community who selected a given subfield label. Note that three other affiliation communities related to neuroscience, human medical science, and veterinary science were also identified in the community analysis of our participant sample (n = 12, 4 and 2). Because these other groups were so small, they are omitted from the summary below.

| | | **Major Affiliation** (% selecting within that clique) | | | |
|---|---|---|---|---|---|
| **Subfield** | % selecting overall | **Eco & Env n = 277 participants** | **Evo & Gene n = 173 participants** | **Biochem & Cell n = 105 participants** | **Physiology n = 39 participants** |
| Biochemistry and Molecular Biology | 15.5 | 0 | 7.5 | 78.1 | 0 |
| Biodiversity and Conservation Biology | 16.3 | 28.5 | 12.1 | 0 | 0 |
| Biophysics | 0.8 | 0 | 0.6 | 2.9 | 2.6 |
| Cell Biology | 4.7 | 0.4 | 1.2 | 23.8 | 2.6 |
| Developmental Biology | 4.2 | 0.4 | 10.4 | 5.7 | 2.6 |
| Ecology | 38.2 | 83.0 | 2.3 | 0 | 0 |
| Environmental Sciences | 10.5 | 20.9 | 2.3 | 1.9 | 0 |
| Evolutionary Biology | 30.9 | 17.0 | 82.1 | 0 | 0 |
| Genetics and Heredity | 6.9 | 2.9 | 16.8 | 4.8 | 0 |
| Medicine (Human) | 1.5 | 0 | 1.7 | 1.9 | 0 |
| Microbiology | 4.2 | 1.4 | 0.6 | 20.0 | 10.3 |
| Neuroscience | 3.3 | 0.7 | 0.6 | 1.0 | 0 |
| Physiology | 9.8 | 1.4 | 7.5 | 4.8 | 97.4 |
| Plant Sciences | 4.9 | 3.6 | 8.1 | 5.7 | 0 |
| Veterinary Medicine | 0.7 | 0.4 | 0.6 | 0 | 0 |

**Table S2. Analysis of the prior heuristic test.** Ordinal logistic regression models used to explain response variation in the prior heuristic test, ranked by model support (n = 623 participants). The best-supported model includes an interaction between the treatment and a participant's level of research experience. See Table S4 and Figures 2 and S2-S3 for further details.

| Rank | Model | Delta AICc | Akaike Weight |
|---|---|---|---|
| 1 | Model 5: response ~ treatment * experience + dataset | 0 | 0.70 |
| 2 | Model 4: response ~ treatment * training + dataset | 3.32 | 0.13 |
| 3 | Model 3: response ~ treatment + experience + dataset | 3.75 | 0.11 |
| 4 | Model 1: response ~ treatment + dataset | 5.54 | 0.04 |
| 5 | Model 2: response ~ treatment + training + dataset | 7.60 | 0.02 |

**Table S3. Analyses of the p-value heuristic test.** Ordinal logistic regression models used to explain response variation in the prior heuristic test, ranked by support (n = 623 participants). The top three models received similar levels of support. See Table S5 and Figures 2 and S2-S3 for further details.

| Rank | Model | Delta AICc | Akaike Weight |
|---|---|---|---|
| 1 | Model 3: response ~ treatment + experience + dataset | 0 | 0.39 |
| 2 | Model 1: response ~ treatment + dataset | 0.86 | 0.25 |
| 3 | Model 5: response ~ treatment * experience + dataset | 1.03 | 0.23 |
| 4 | Model 2: response ~ treatment + training + dataset | 2.91 | 0.09 |
| 5 | Model 4: response ~ treatment * training + dataset | 4.40 | 0.04 |

**Table S4. Best-supported model for the prior heuristic test.** Coefficients estimates are reported for the best-supported ordinal logistic regression model from Table S2 (n = 623 responses). The model also includes a categorical predictor for the scatterplot dataset shown, with 6 levels; these dataset coefficients are omitted here for simplicity. See Figures 2 and S2-S3 for further details.

| Model 5 | Estimate | SE | z | p-value |
|---|---|---|---|---|
| treatment | 1.35 | 0.31 | 4.39 | < 0.0001 |
| experience | 0.03 | 0.10 | 0.30 | 0.76 |
| treatment:experience | –0.33 | 0.14 | –2.40 | 0.02 |

**Table S5. Best-supported models for the p-value heuristic test.** Models 3, 1 and 2 received similar support in this analysis as shown in Table S3. As such, coefficient estimates are reported below for all three ordinal models (n = 623 responses). Each model also includes a categorical predictor for the scatterplot dataset shown, with 6 levels; these dataset coefficients are omitted here for simplicity. See Figures 2 and S2-S3 for further details.

| Model 3 | Estimate | SE | z | p-value |
|---|---|---|---|---|
| treatment | 0.70 | 0.15 | 4.50 | < 0.0001 |
| experience | –0.11 | 0.07 | –1.71 | 0.09 |

| Model 1 | Estimate | SE | z | p-value |
|---|---|---|---|---|
| treatment | 0.71 | 0.15 | 4.57 | < 0.0001 |

| Model 5 | Estimate | SE | z | p-value |
|---|---|---|---|---|
| treatment | 0.95 | 0.29 | 3.23 | 0.001 |
| experience | –0.05 | 0.09 | –0.57 | 0.57 |
| treatment:experience | –0.13 | 0.13 | –1.02 | 0.31 |

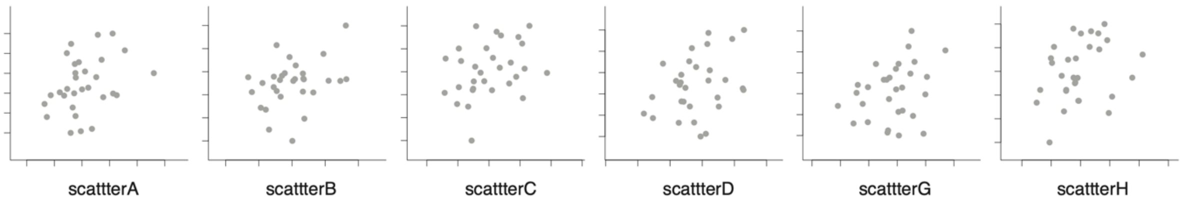


**Figure S1. Dataset scatterplots used to test data interpretation heuristics.** Each scatterplot presents 30 datapoints with an ambiguous, positive correlation (Pearson's R ranging 0.29 – 0.24; true p-values ranging from 0.07 to 0.11). Each participant viewed two different scatterplots in the survey (one for each heuristic test), and scatterplots were randomized across manipulations within the two tests.

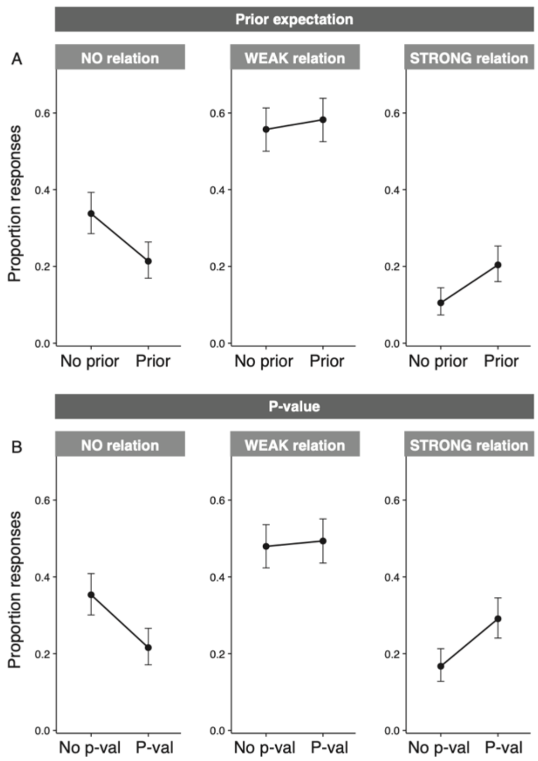


**Figure S2. Main effects of the prior and p-value heuristics on data interpretation**. Columns show the proportion of participants who interpreted that there was “No relationship” (left), “A weak positive relationship” (middle), or “A strong positive relationship” (right). Both manipulations, (A) the prior and (B) the p-value, affected data interpretations. While Figure 2 stratifies participants by experience level, here we show values for the entire participant pool. Error bars show 95% confidence intervals.

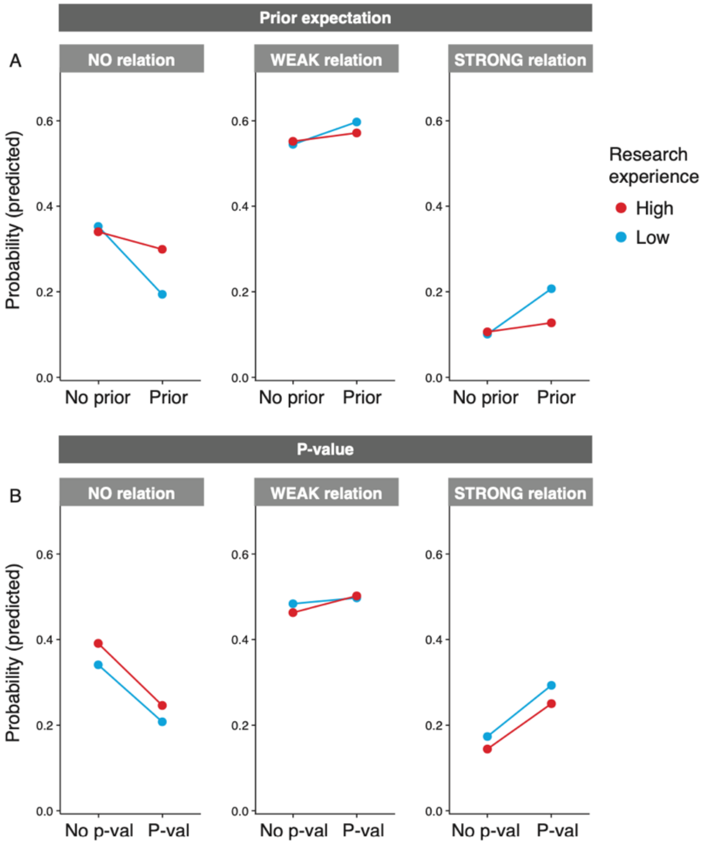


**Figure S3. Best-supported models of data interpretation responses.** The layout of this figure follows Figure 2 of the main text which shows the proportion of survey responses. This figure shows the predictions of the best-supported models from Table S4 and S5.

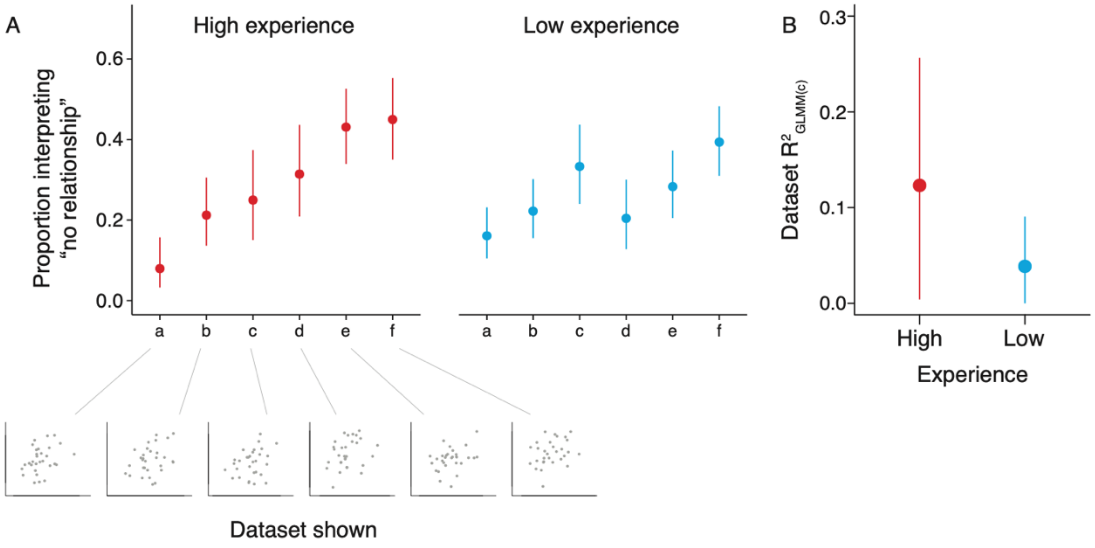


**Figure S4. Are highly experienced researchers more sensitive to the data at hand?** To investigate this question, we stratified researchers into two experience groups, low research experience (0-5 recent peer-reviewed research articles) and high research experience (6 or more recent peer-reviewed research articles). We then investigated the extent to which response variation could be explained by the datasets shown. (A) Responses for the six datasets, pooled across the two heuristic tests. Some of the ambiguous datasets were viewed as more likely to represent a relationship, whereas others were viewed more often as having no relationship. (B) $R^2$ estimates for the proportion of response variance explained by dataset. In the high-experience group, differences among datasets explained slightly more of the response variation, as compared to the low-experience group. This suggests that more experienced researchers may be more sensitive to the data at hand. Error bars show 95% confidence intervals.

***Screenshots of the survey***

### Data Visualization Survey

**Participation in this study is voluntary and will take 1-2 minutes.**
The aim of this study is to investigate how scientists interpret data visualizations. This study is a brief multiple-choice survey that takes 1-2 minutes to complete. Your participation in this survey is voluntary, and you may choose not to take part. There are no risks associated with participating in this survey.

**If you complete the survey, you can choose to enter an optional draw to win a $40 or 40£ Amazon gift card (one entry per participant).** You may enter the draw after completing the survey by providing your email address. The winner will be selected randomly when the survey closes on March 31, 2021, and will be notified of their gift card by email. The odds of winning the draw are not known in advance and will depend on the number of entrants. After completion of the draw on March 31, all email addresses will be deleted.

Your data will be stored and protected by Qualtrics on a server located in Toronto, Canada, but may be disclosed via a court order or data breach. Only the survey responses will be stored as confidential data by the researchers. These responses will be analyzed for peer-reviewed research on the interpretation of data. The responses will be published in anonymized form (i.e., without any identifying information) for reproducibility and use in future research.

**By following the link below to complete this survey, you are agreeing to participate in the study.**
Should you choose to participate, you may withdraw from the study up until the point of submission. You can withdraw prior to submission by exiting the survey or closing the browser window.

This research has been cleared by Carleton University Research Ethics Board-B (CUREB-B Clearance #114980). Should you have any ethical concerns with this study, please contact the Carleton University Research Ethics Board. For all other questions about the research, please contact us at the email below.

Roslyn Dakin and Ashley Irwin
Department of Biology
Carleton University

Funding supported by Carleton University

→

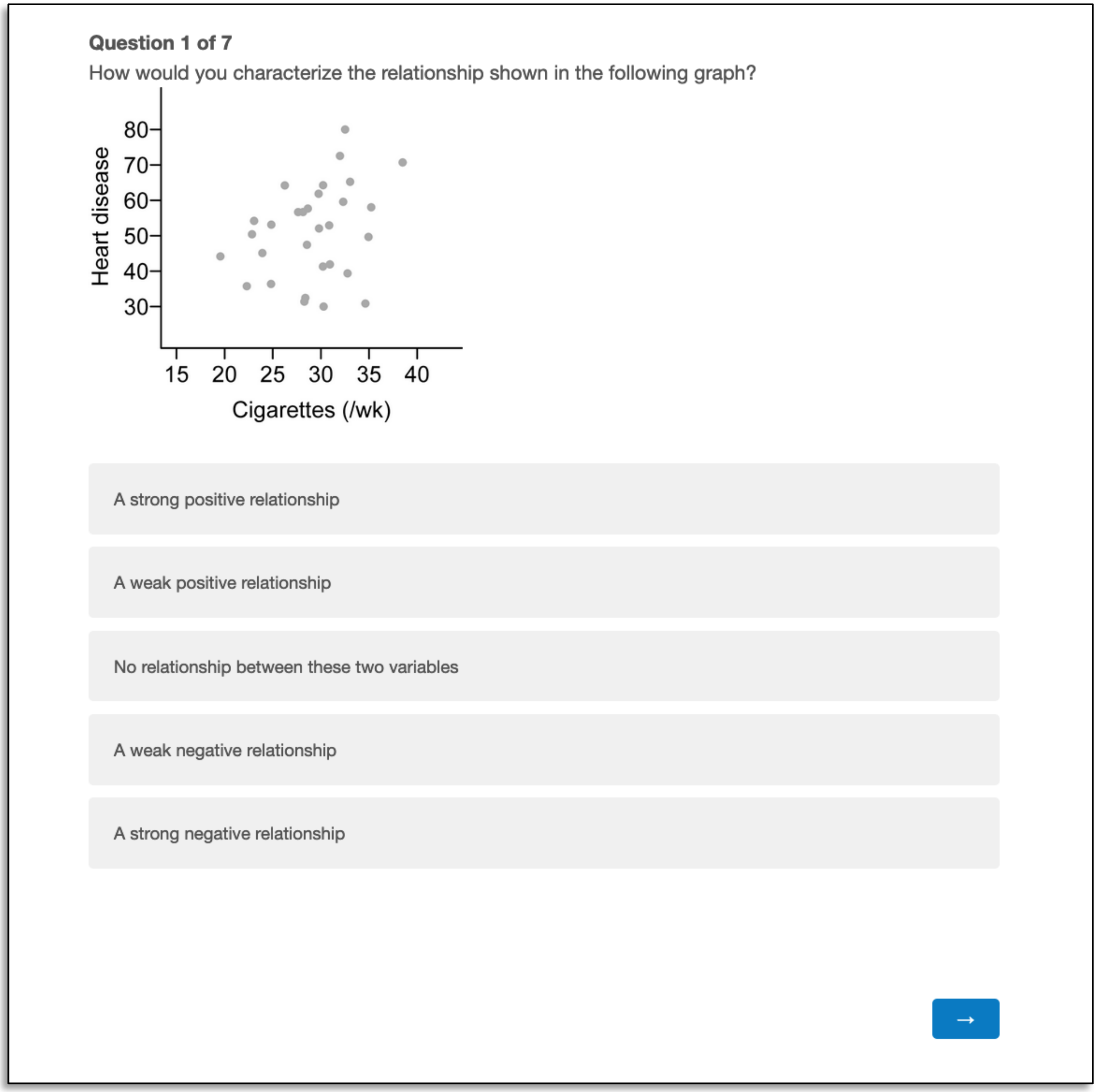


** Note that some participants saw a version of this question with the axes labelled "Internet browsing" and "Height (/in.)", as shown in Figure 1 of the main text. This question tests the prior heuristic. Note also that different participants saw other ambiguous scatterplots (see Figures S1 and S4).*

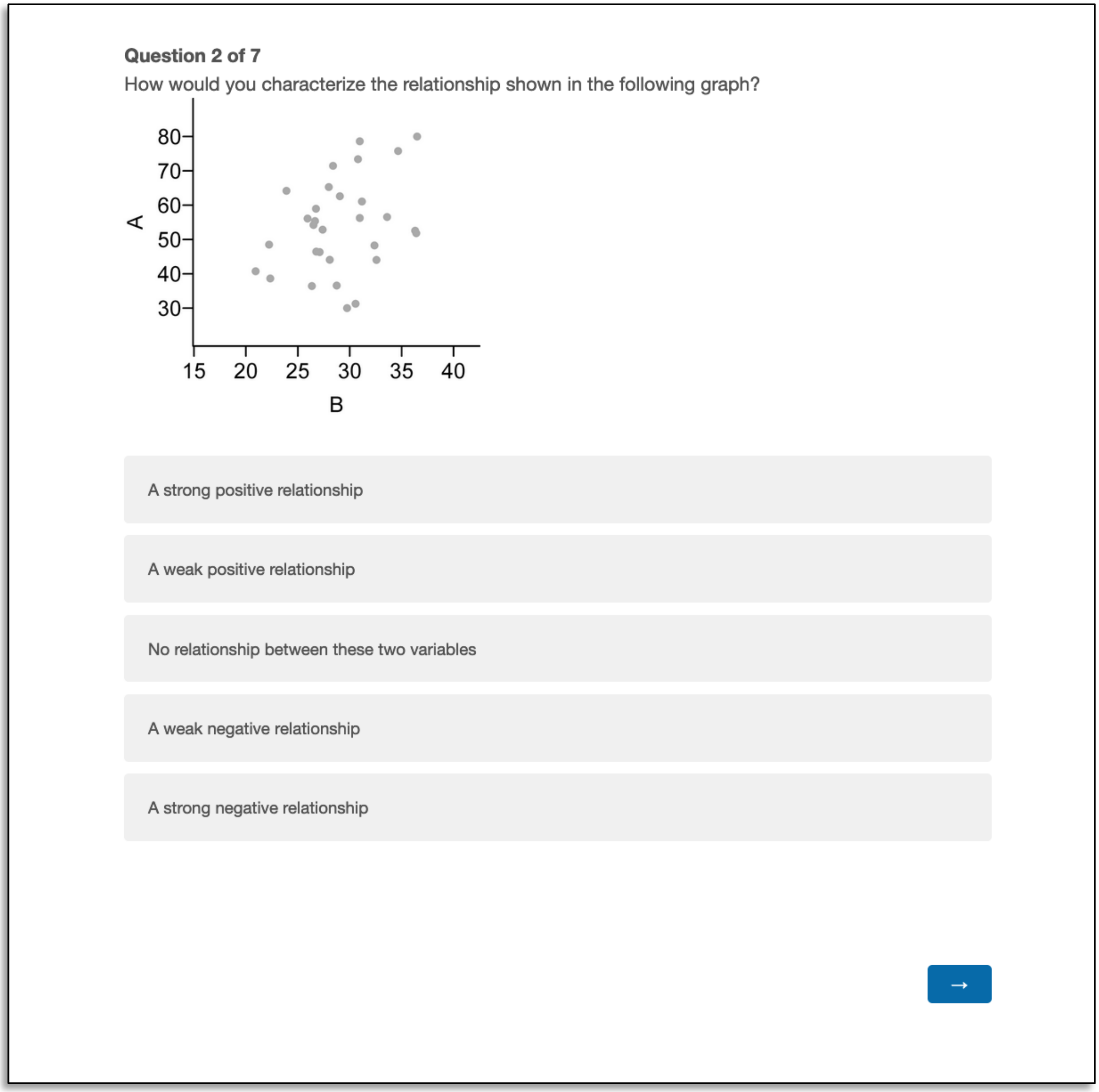


*** Note that some participants saw a version of this question with text within the graph panel that says " *** p < 0.0001", as shown in Figure 1 of the main text. This question tests the p-value heuristic. Note also that different participants saw other ambiguous scatterplots (see Figures S1 and S4).*

**Question 3 of 7**
What is your highest degree obtained?

- Postgraduate degree (e.g., MSc, PhD, MD, DVM, or similar )
- Some postgraduate degree (i.e., in progress)
- Bachelor's or undergraduate degree
- Other

**Question 4 of 7**
Approximately how many courses in statistics have you completed (as a student) or taught (as an instructor)? (Include undergraduate and graduate level)

0
1
2
3
4
5
6
7
8
9 or more

**Question 5 of 7**
How many peer-reviewed research articles have you published in the last 5 years?

- 0-5
- 6-10
- 11-15
- 16 or more

**Question 6 of 7**
Please select the country of your primary university:

- Australia
- Canada
- England
- United States
- Other

118

**Question 7 of 7**
Please select one or more field(s) that you identify most closely with:

- Biochemistry and Molecular Biology
- Biodiversity and Conservation Biology
- Biophysics
- Cell Biology
- Developmental Biology
- Ecology
- Environmental Sciences
- Evolutionary Biology
- Genetics and Heredity
- Medicine (Human)
- Microbiology
- Neuroscience
- Physiology
- Plant Sciences
- Veterinary Medicine
- Other (please describe)

If you wish to enter the optional draw for a $40/40£ gift card, please type your email address below. If you do not wish to enter the draw, leave this space blank. This email address will be used to contact the winning entry.

Email ____________________

**Click the Button Below to Submit Your Survey**

Thank you for your participation!

Submit

120

We thank you for your time spent taking this survey.
Your response has been recorded.

122
